# Reference genes for qRT-PCR normalisation in different tissues, developmental stages and stress conditions of *Hypericum perforatum*

**DOI:** 10.1101/499186

**Authors:** Wen Zhou, Lei Yang, Yan Sun, Qian Zhang, Bin Li, Bin Wang, Lin Li, Donghao Wang, Shiqiang Wang, Zhezhi Wang

## Abstract

*Hypericum perforatum* is a widely known medicinal herb used mostly as a remedy for depression because of its abundant secondary metabolites. Quantitative real-time PCR (qRT-PCR) is an optimized method for the efficient and reliable quantification of gene expression studies. In general, reference genes are used in qRT-PCR analysis because of their known or suspected housekeeping roles. However, their expression level cannot be assumed to remain stable under all possible experimental conditions. Thus, the identification of high quality reference genes is very necessary for the interpretation of qRT-PCR data. In this study, we investigated the expression of fourteen candidate genes, including nine housekeeping genes and five potential candidate genes. Additionally, the *HpHYP1* gene, belonging to the PR-10 family associated with stress control, was used for validation of the candidate reference genes. Three programs were applied to evaluate the gene expression stability across four different plant tissues, three developmental stages and a set of abiotic stress and hormonal treatments. The candidate genes showed a wide range of Ct values in all samples, indicating that they are differentially expressed. Integrating all of the algorithms and evaluations, *ACT2* and *TUB-β* were the most stable combination overall and for different developmental stages samples. Moreover, *ACT2* and *EF1-α* were considered to be the two most applicable reference genes for different tissues and for stress samples. Majority of the conventional housekeeping genes exhibited better than the potential reference genes. The obtained results will contribute to improving credibility of standardization and quantification of transcription levels in future expression research of *H. perforatum*.

## Introduction

An increasing number of studies on the gene expression levels in plants have been carried out in order to understand the signalling and metabolic pathways underlying the developmental processes involved in plant development and growth, as well as in plant responses to biotic and abiotic stresses [1–4]. Methods for detecting gene expression include northern blot, gene chips, semi-PCR, RNase protection analysis and qRT-PCR. qRT-PCR because of its high sensitivity, specificity, reproducibility and accuracy has become a very popular and effective method for detecting and quantifying gene transcription levels [5–7]. Reliable quantification of gene expression levels by qRT-PCR analysis requires the standardization and fine-tuning of several parameters, such as the amount of initial sample, RNA recovery and integrity, enzymatic efficiency of cDNA synthesis and PCR amplification, and the overall transcriptional activity of the tissues or cells analysed [5–9]. The expression stability of frequently used reference genes also cannot be neglected. Therefore, normalization for transcript levels of test genes is essential for minimizing technical differences for different samples and experimental conditions [9]. More importantly, selection of stable reference gene(s) is prerequisite before qRT-PCR analysis.

A appropriate reference gene is considered to be not affected by the experimental conditions and also shouldh maintained at a invariable levle among samples [10]. Actually, the transcription levels of the traditional reference genes and even some housekeeping genes (HKGs) can vary between different types of tissue and under different treatment conditions [11]. Therefore, a growing number of research have been published on the analysis and evaluation of the stability of internal reference genes in plant tissues under different conditions. For *Gentiana macrophylla, SAND1* and *EF-1α4* were found to be the most suitable overall; *GAPC2* and *SAND1* certified to be the best reference genes for the roots under abiotic stresses, while *SAND1* and *EF-1α4* were screened out from the leaves of stressed plants [12]. For the tung tree, *ACT7, UBQ, GAPDH*, and *EF-1α* were the four optimal reference genes in all samples and developing seeds. *ACT7, EF-1β, GAPDH*, and *TEF1* were the top four candidate genes for different tissues [13]. Consequently, different reference genes should be applied in different experimental materials and conditions. In addition, the reference genes with lower stability may lead to a more erroneous understanding of qRT-PCR [14] and mask the true nature of gene expression [15,16].

*Hypericum perforatum* L. (commonly known as St. John’s wort) is a widely known medicinal herb used mostly as a remedy for depression [17]. Pure compounds essentially isolated from *H. perforatum*, namely, naphthodianthrones and phloroglucinols, are proven to possess anti-depressive, anti-cancer, anti-viral, anti-inflammatory activities and other activities [18,19]. Xanthones and flavonoids have also been identified in extracts from this plan [20]. To date, a limited number qRT-PCR studies focusing on *H. perforatum* has been published. Isabel et al. studied the stability of 11 candidate reference genes analysed in *H. perforatum* plants only subjected to cold and heat stresses, revealing TUB being the most stable gene in both experimental conditions [21]. Therefore, it is very necessary to screen suitable reference genes in different tissues of *H. perforatum* under different experimental conditions. Ribosomal RNA and some HKGs usually served as reference genes, such as actin (*ACT*), tubulin (*TUB*), glyceraldehyde-3-phosphate dehydrogenase (*GAPDH*) and polyubiquitin (*UBQ*) [22–26], whereas many studies have revealed that the most commonly used HKGs are not always reliable in different experimental samples [27–30]. So a evaluation for screening out the optimal HKGs in different species is essential.

This study aimed to assess the expression stabilities of fourteen reference genes in fifteen experimental samples by qRT-PCR, including 9 traditional HKGs and 5 potential reference genes: *GAPDH*, actin (ACT2, ACT3, *ACT7*), ubiquitin-conjugating (*UBC2*), elongation factor (*EF-1a*), tubulin (*TUB-α, TUB-β*), cyclophilin (*CYP1*), polyketide synthase (*PKS1*), glutamate semialdehyde aminomutase (*GSA*), SAND family protein (*SAND*), ribosomal protein L (*RPL13*) and protein phosphatase 2A (*PP2A*). These genes were selected from the *H. perforatum* genome sequencing data in our lab. Three mathematical programs, geNorm [31], NormFinder [32] and BestKeeper [33], were applied to analyse the raw Ct values and to figure out the relatively stable expressed genes. All of the raw Ct values need to be converted to ΔCt (ΔCt = each corresponding average Ct value - mimimum Ct value) in NormFinder and geNorm algorithms, while there is no need in BestKeeper [34]. The selected reference genes were detected by using a target gene *HpHYP1* belonging to the Pathogenesis related class-10 (PR-10) family [35,36].

## Materials and Methods

### Plant materials

The *H. perforatum* seeds (2n=2x=16) were germinated on a seedling bed in the glasshouse (25 ± 2 °C, natural lighting, 60–80% humidity). Whole plant tissues were collected at the one-month-old (1M), two-month-old (2M), three-month-old (3M) and six-month-old (6M) stages. The different tissue samples (leaf, flower, stem and root) were taken from two-year-old plants (2n=2x=16). Three-month-old seedlings were treated with abiotic stress and hormonal treatments, including 10 μM salicylic acid (SA), 200 μM methyl jasmonate (MeJA), 100 μM abscisic acid (ABA), 1 mM AgNO3 (Ag), 200 μM CuSO_4_ (Cu), 100 mM NaCl (Na), low temperature (4 °C) and wounding (W). The stress samples were acquired severally after 6 h of the corresponding treatments, and the control groups were collected following non-treatment. The collected plant samples are. All samples were collected in three replicates and frozen in liquid nitrogen immediately and then stored at −80 °C.

### Total RNA isolation and cDNA synthesis

Total RNA was extracted using the Polysaccharide and Polyphenols Plant Quick RNA Isolation Kit (centrifugal column type; Waryong, Beijing, China). The genomic DNA was digested with RNase-free DNase I (TaKaRa, Japan). The total RNA was quantified by absorbance at A_260_/A_280_ nm and A_260_/A_230_ nm with a NanoDrop 2000c spectrophotometer (Thermo Scientific, USA). A 1% (p/v) agarose gel was run to visualize the integrity of the RNA. Only RNA samples with a A_260_/A_280_ wavelength ratio between 1.9 and 2.1 and an A_260_/A_230_ ratio close to 2.0 were used for cDNA synthesis. A quantity of 1.0 μg DNA-free total RNA was used to synthesize first-strand cDNAs with a PrimeScript RT Reagent Kit (TaKaRa, China) in a 20 μL volume. All cDNA samples were diluted (1:30) with DNase/RNase-free deionized water for qRT-PCR.

### Selection of reference genes and primer design

We performed genomic sequencing of *H. perforatum* using Illumina paired-end, 10X Genomics linked reads and PacBio SMART (unpublished). The transcriptome sequencing of *H. perforatum* for the roots, stems, leaves and flowers assisted annotation. Fourteen candidate genes, including 9 traditional HKGs (*GAPDH, ACT2, ACT3, ACT7, EF1-a UBC2, TUB-α, TUB-β* and *CYP1*) and 5 potential reference genes (*PKS1, GSA, RPL13, SAND and PP2A*), were selected for investigation in order to identify the most stably expressed reference genes. To ensure the accuracy of the reference gene predictions, we first screened the candidate genes according to the genome annotation of each, which was assigned based on the best match of the alignments using Blastp to SwissProt, KEGG, NR and TrEMBL databases. Then the coding sequences of the fourteen chosen genes were used as queries for BLAST orderly through the TAIR ((http://www.arabidopsis.org/) for further ensuring accuracy. Finally the sequences with the highest homology with Arabidopsis were shown in Table 1. The primers of all the genes were designed using GenScript (https://www.genscript.com) with a melting temperature between 59 and 61 °C, a primer length of 20 - 25 bp, and an amplicon length of 70 - 180 bp. The descriptions of the candidate reference genes, primer sequences and qRT-PCR amplification efficiencies are presented in Table 1.

### qRT-PCR conditions and analysis

PCR reactions were performed on the Roche LightCycler 96 system using SYBR® Master Mix. Reactions were done in 20 μl volumes in triplicate containing 5 μL of 30-fold diluted synthesized cDNA, 10 μL SYBR® Master Mix, 0.4 μL 10 mM forward primer, 0.4 μL 10 mM reverse primer and 4.2 μL DNase/RNase-free deionized water. The cycling conditions included 95 °C for 30 s, 45 cycles of 95 °C for 5 s and 60 °C for 30 s, followed by a final melting curve analysis. Each reaction was set up with a negative control group and conducted in three technical replicates. According to calculate the amplification efficiency from 10-fold continuous dilution of the cDNA for each gene, the standard curves were constructed to get the correlation coefficients (R^2^) and slope values. So the corresponding PCR amplification efficiencies (E) could be calculated (E = (10^− 1/slope^ − 1) × 100 [37]).

### Assessment of expression stability

The expression stability of the genes was analysed with three different Visual Basic applets, GeNorm [30,38], NormFinder [39, 40] and BestKeeper [41,42]. GeNorm derives a stability measure (M-value), via a stepwise exclusion of the least stable gene, and creates a stability ranking [43]. Genes with M < 1.5 are generally considered stable reference genes [44,45]. This measure is based on the principle that the expression ratio of two ideal control genes should be identical in all samples; thus genes with the lowest M-value are the most stably expressed [46]. NormFinder uses an ANOVA-based model to estimate intra- and inter-group variation within tissues or treatments (“groups” in NormFinder terminology) to assess the expression stability [47]. For GeNorm and NormFinder, the raw Ct values needed to be converted to relative quantities (Q) by the formula Q = 2^− ΔCt^, in which ΔCt = each average Ct value - mimimum Ct value [48]. In BestKeeper, the coefficient of variance (CV) and the standard deviation (SD) were caculated by the Ct values, with the lower CV and SD indicating higher stability [49]. All other statistical analyses were calculated by GraphPad Prism 6.0 [50], and the level of statistical significance was assessed using * P < 0.05, ** P < 0.01, and *** P < 0.001.

### Validation of reference gene stability

Previous studies showed that the *HpHYP1* (JF774163) gene is classified as a member of the plant PR-10 (pathogenesis-related class 10) gene family. Members of the PR-10 family are ubiquitous plant proteins associated with stress control [51]. The relative expression of *HpHYP1* gene in different tissues were measured and standardized by using the most stable and unstable candidate genes as internal reference to verify the reliability of the selected genes according to the 2^− ΔΔCt^ method [52]. Three technical replicates were performed for each biological sample.

## Results

### Primer specificity and expression level analysis of candidate reference genes

The gene names and abbreviations, accession numbers, primer sequences, amplification efficiencies, amplicon sizes, Tm values and molecular functions are listed in Table 1. The amplification efficiencies ranged from 92.5% (*RPL13*) to 109.5% (*TUB-α*), the Tm values varied from 81.2 °C (*PP2A*) to 87.8 °C (*CYP1*), and the amplicon sizes were between 76 bp (*GAPDH* and *PKS1*) and 170 bp (*TUB-β*). Furthermore, primer specificities were detected by melting curves (Fig 1) and a single band indicated the correct size of each pair. The raw CT values of different genes ranged from 21.2 to 38.2 (Fig 2). Data were analysed within experiments and were divided into four groups: tissues from two-year-old plants (TS: R, S, L, and F), developmental stages seedlings (SG: 1M, 2M, 3M, and 6M), three-month-old seedlings exposed to abiotic stresses (ST: SA, MeJA, ABA, Cu, Ag, Na, 4 °C and W) and a combination of all experimental conditions (TT). Thefourteen selected genes’ expression levels in the four groups were shown in Fig. 2. Among them, UBC2 (SG) performed the highest Ct value (30.33), but GAPDH (TS) represented the opposite, which indicated the levels of their expression.

**Table 1.**
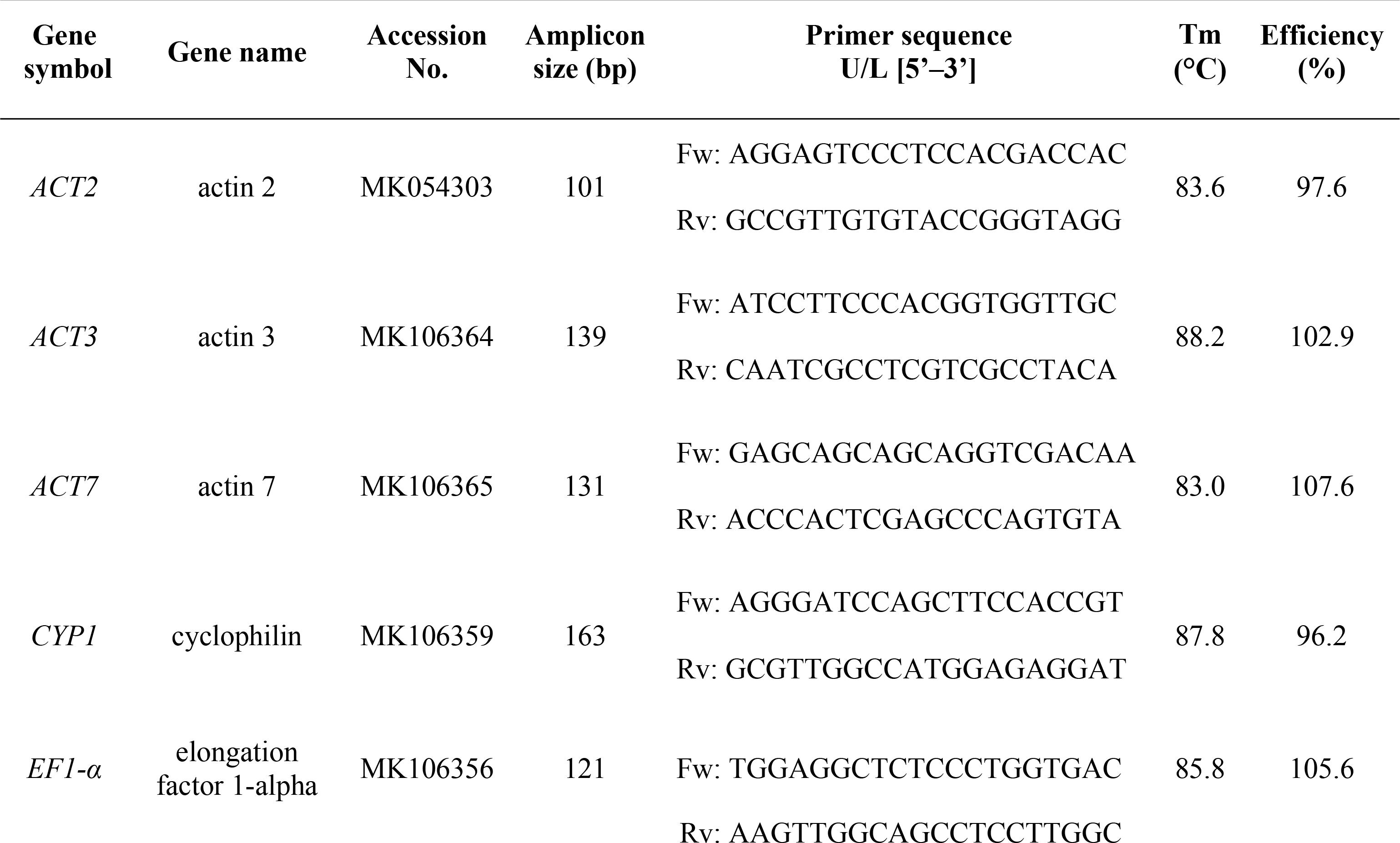
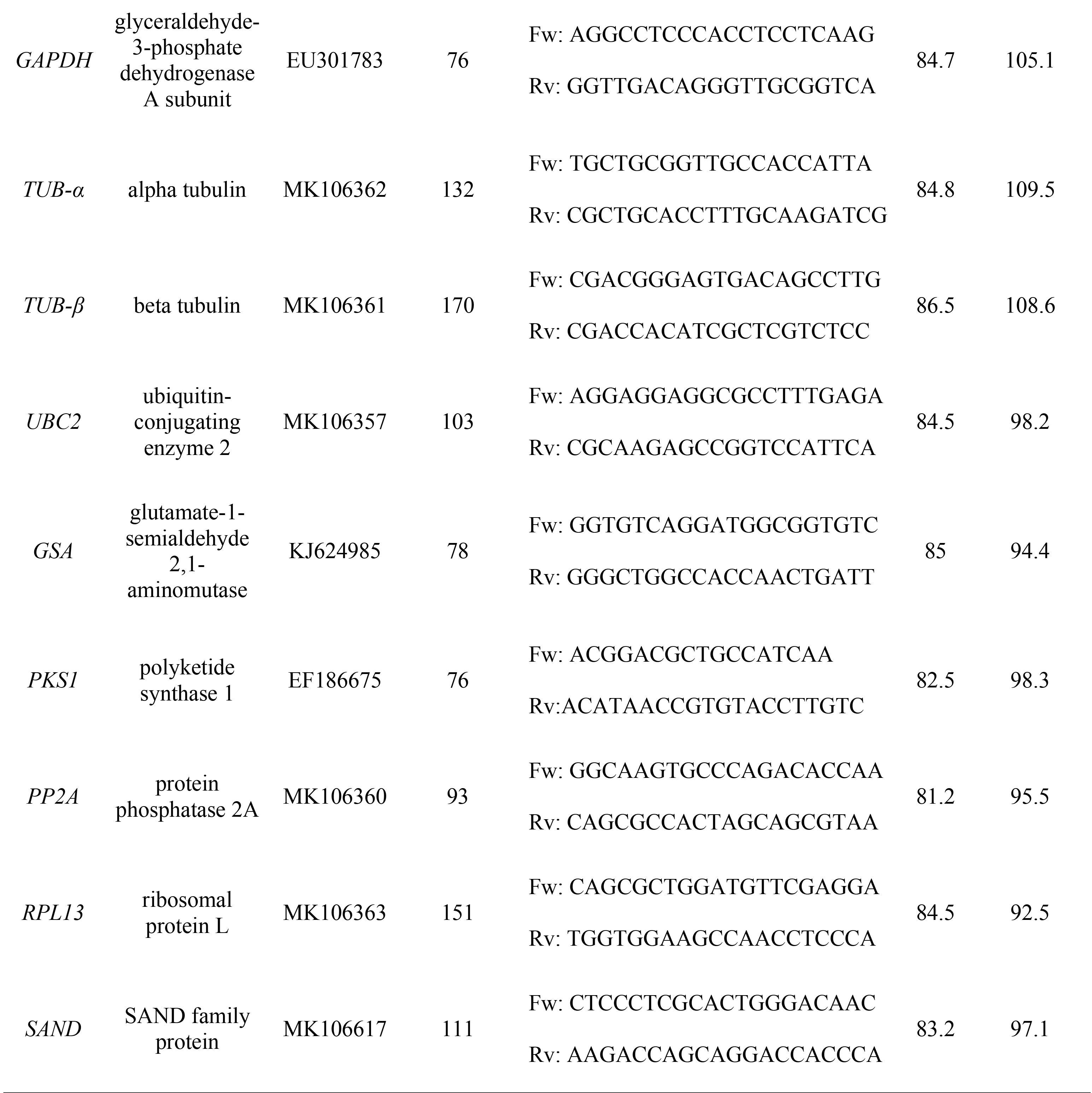
Description of the candidate reference genes, primer sequences and qRT-PCR amplification efficiencies in *H. perforatum*.

**Figure 1.**
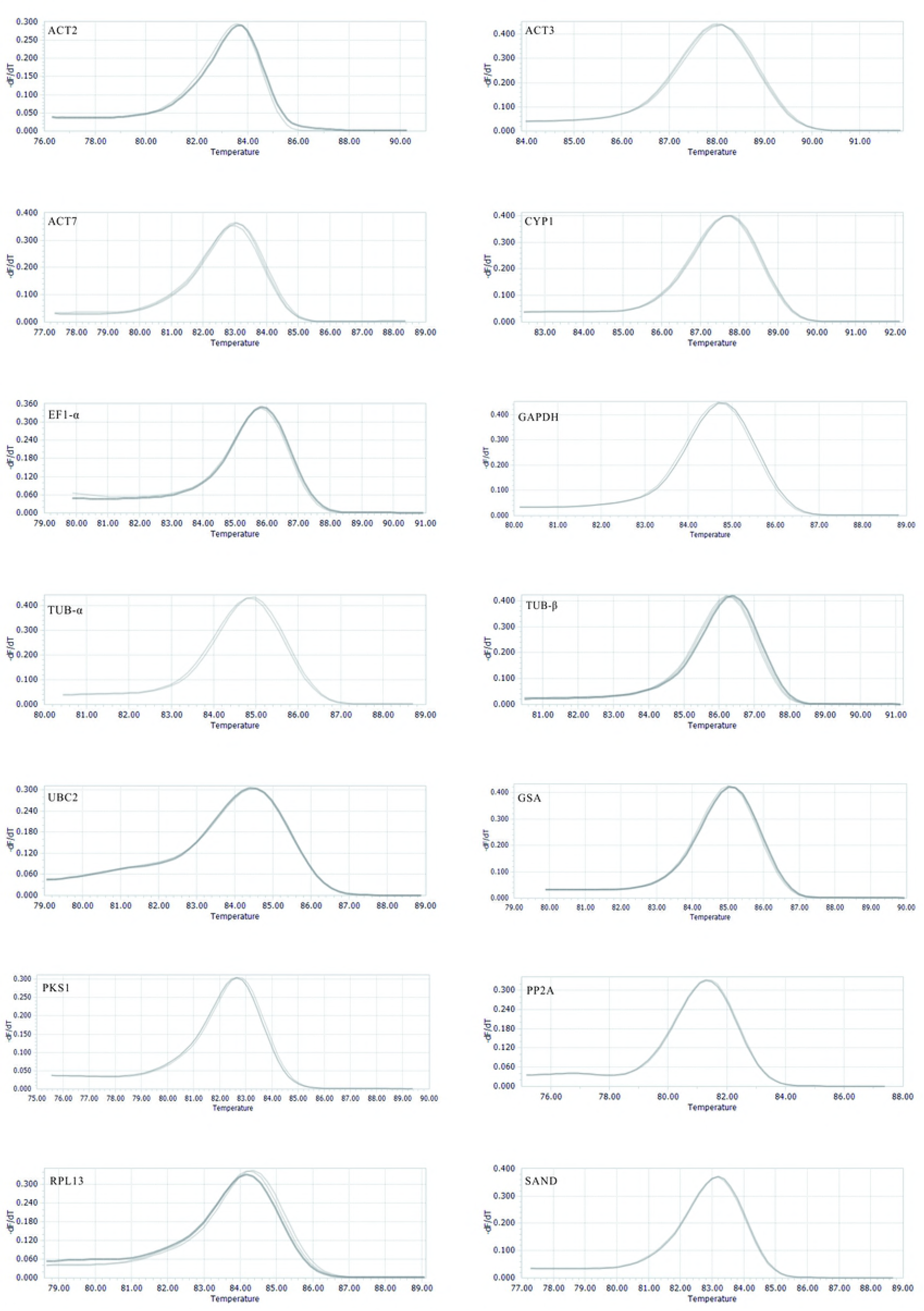
Melting curves for the candidate reference genes. The melting temperature was shown by plotting the negative derivative of fluorescence relative to Celsius temperature (-(d/dT)).

**Figure 2.**
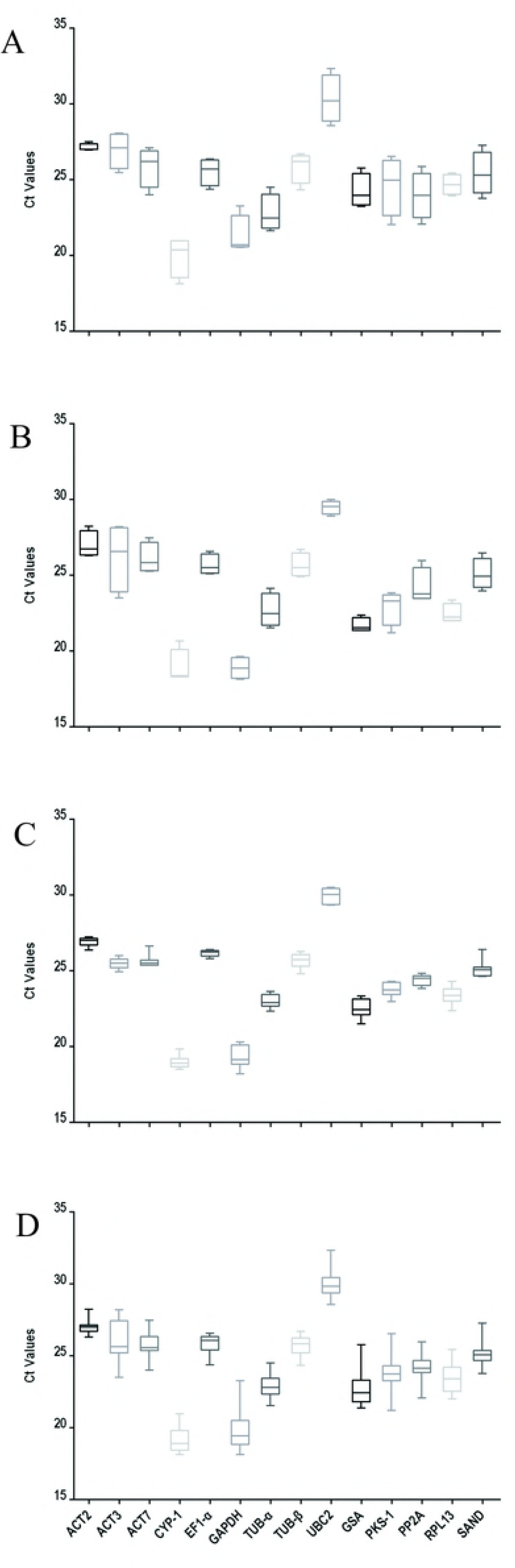
Average Ct values for the candidate reference genes. (A) Different tissues; (B) developmental stage seedlings; (C) stress-treated seedlings; (D) data from all experimental conditions combined. A line across the box is depicted as the median. The box indicates the 25th and 75th percentiles; whisker caps represent the maximum and minimum values; and dots represent outliers.

### Stability of candidate reference genes

GeNorm is requested to caculate a normalization factor from the geometric mean of the genes to distinguished the most stably expressed gene. The M-values is defined as the average pairwise variation of a particular gene with all other potential reference genes. In GeNorm, the threshold for eliminating a gene was set as *M* < 1.5, the lower the M value, the higher the stability [53]. The expression stability ranking of the fourteen reference genes was arranged in the TT samples as follows: *TUB-β* > *ACT2* > *CYP1* > *EF1-α* > *TUB-α* > *SAND* > *UBC2* > *ACT7* > *PP2A* > *GSA* > *PKS1* > *RPL13* > *GAPDH* > *ACT3. ACT2* and *ACT3* were the genes with the lowest and highest M values in the TS, SG and ST groups. In brief, *ACT2* and *TUB-β* were the optimal two reference genes for pooled samples that included all treatments. *ACT7, RPL13* and *GAPDH* were determined to be the relatively unstable reference genes in the majority of samples (Table 2, Fig 3). The Genorm pairwise variation (V) values between ranked genes (V_n_/V_n+1_) were determined for the optimal number of reference genes to be used in the quantitative analysis experiments. A cut-off of 0.15 (V_n_ value) is usually applied [53]. The V_2/3_ values for all of the experimental sets in *H. perforatum* were lower than the cut off threshold of 0.15 (Fig 4), which indicated that the combination of two reference genes can accurately standardize these samples. In other words, *ACT2* and *TUB-β* was the optimum selection as the reference gene combination for qRT-PCR analysis in all of the experimental sets.

**Table 2.** Ranking and M-values of the candidate reference genes calculated by geNorm.

| Rank | Total |  | Tissue |  | Stage |  | Stress |  |
| --- | --- | --- | --- | --- | --- | --- | --- | --- |
|  | Gene | M-value | Gene | M-value | Gene | M-value | Gene | M-value |
| 1 | *TUB-β* | 0.818 | *ACT2* | 0.861 | *ACT2* | 0.639 | *ACT2* | 0.474 |
| 2 | *ACT2* | 0.890 | *EF1-α* | 0.867 | *TUB-β* | 0.665 | *EF1-α* | 0.501 |
| 3 | *CYP1* | 0.895 | *TUB-β* | 0.882 | *CYP1* | 0.670 | *CYP1* | 0.504 |
| 4 | *EF1-α* | 0.936 | *CYP1* | 0.921 | *TUB-α* | 0.737 | *ACT7* | 0.506 |
| 5 | *TUB-α* | 0.946 | *TUB-α* | 0.934 | *GSA* | 0.738 | *TUB-β* | 0.506 |
| 6 | *SAND* | 0.952 | *ACT7* | 1.009 | *EF1-α* | 0.748 | *TUB-α* | 0.531 |
| 7 | *UBC2* | 1.005 | *UBC2* | 1.070 | *PP2A* | 0.750 | *GSA* | 0.559 |
| 8 | *ACT7* | 1.009 | *PP2A* | 1.126 | *GAPDH* | 0.753 | *PKS1* | 0.599 |
| 9 | *PP2A* | 1.059 | *SAND* | 1.167 | *SAND* | 0.762 | *SAND* | 0.639 |
| 10 | *GSA* | 1.086 | *GSA* | 1.181 | *UBC2* | 0.862 | *UBC2* | 0.642 |
| 11 | *PKSI* | 1.087 | *PKSI* | 1.199 | *ACT7* | 0.886 | *GAPDH* | 0.645 |
| 12 | *RPL13* | 1.089 | *RPL13* | 1.229 | *PKSI* | 0.940 | *PP2A* | 0.650 |
| 13 | *GAPDH* | 1.151 | *GAPDH* | 1.339 | *RPL13* | 1.205 | *RPL13* | 0.700 |
| 14 | *ACT3* | 1.216 | *ACT3* | 1.404 | *ACT3* | 1.284 | *ACT3* | 0.772 |

**Figure 3.**
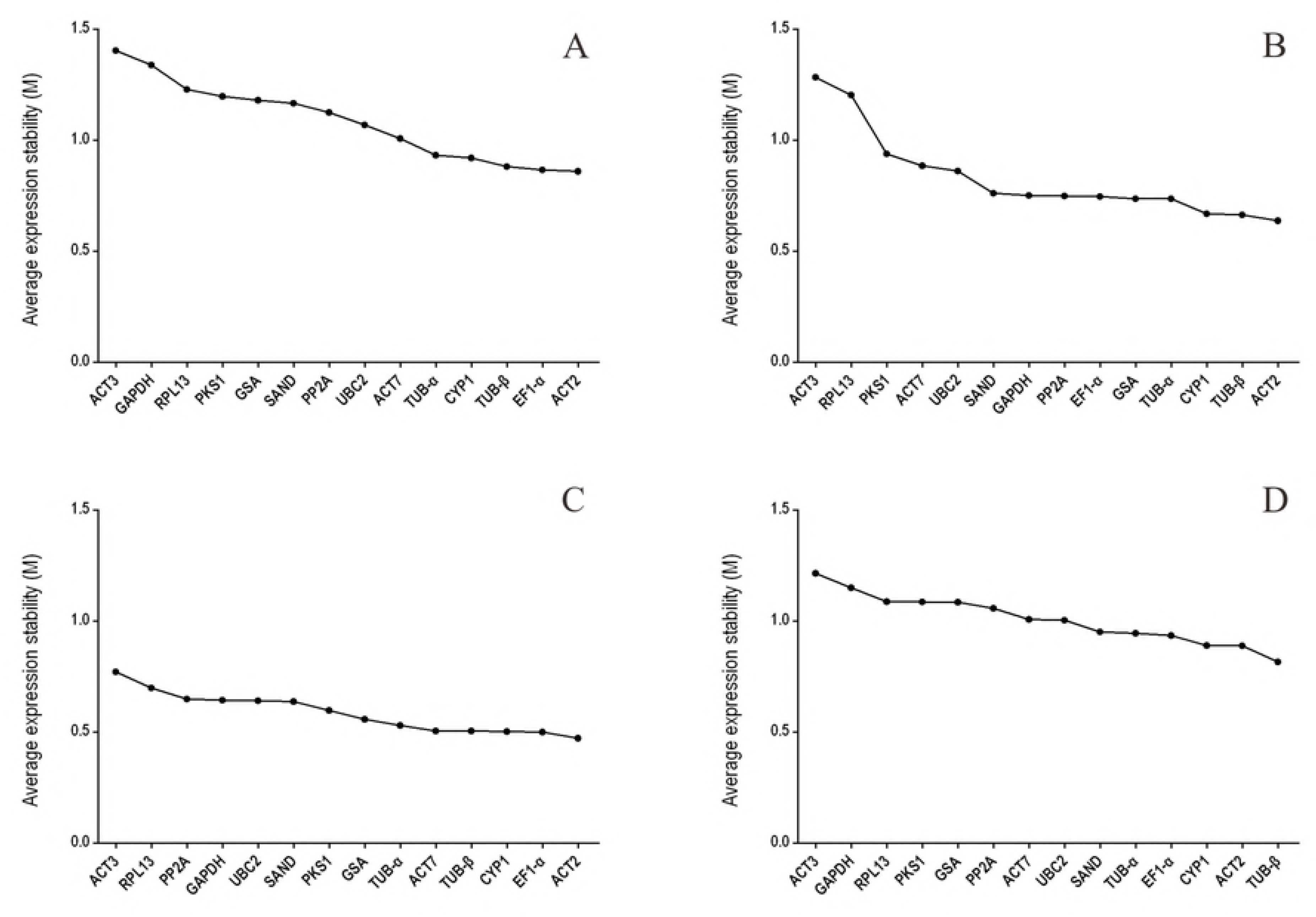
Average expression stability (M-value) of 14 candidate reference genes calculated by geNorm. (A) Different tissues; (B) developmental stage seedlings; (C) stress-treated seedlings; (D) data from all experimental groups combined. A lower average expression stability (M-value) indicates more stable expression.

**Figure 4.**
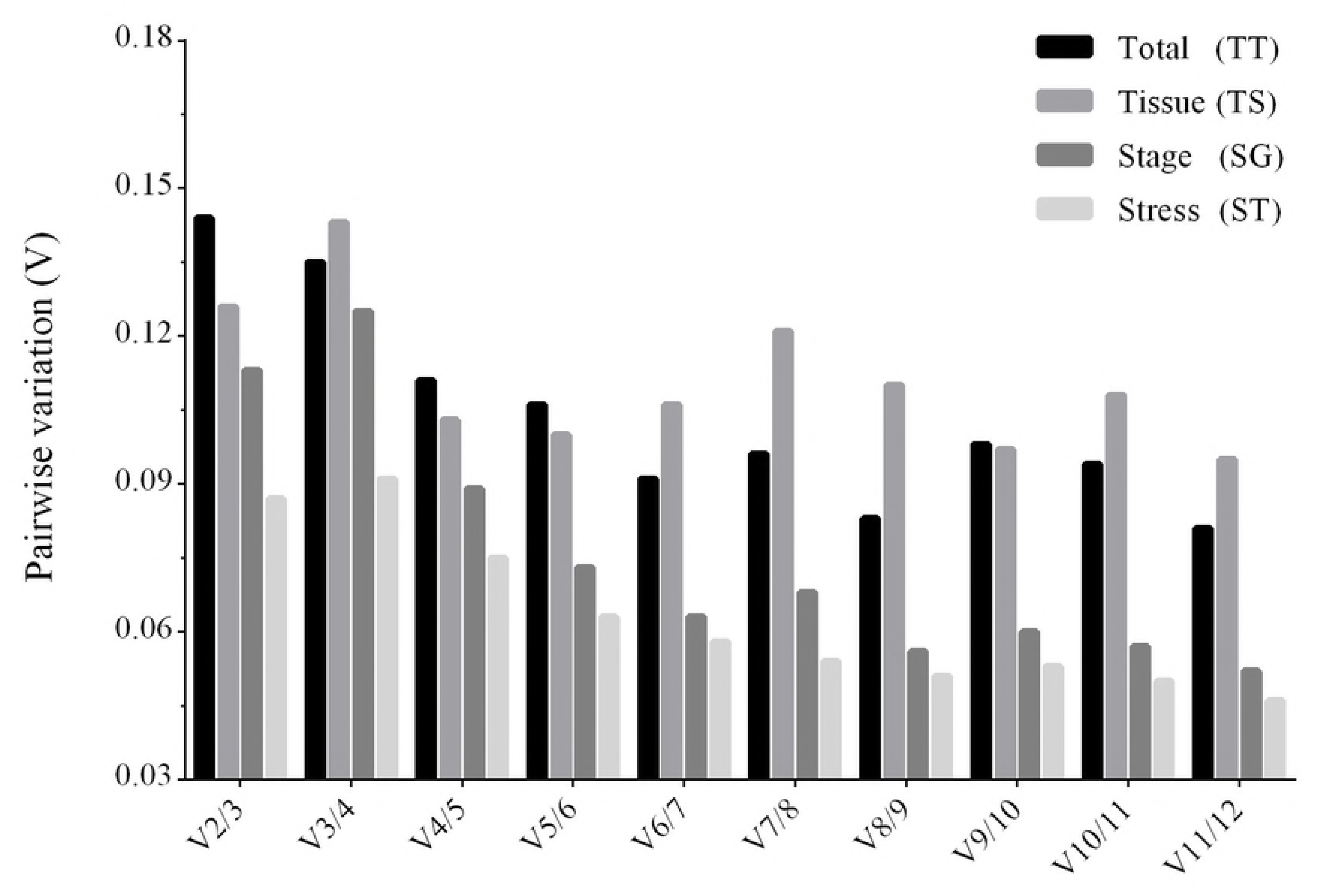
Pairwise variation (V) calculated by geNorm to determine the optimal number of reference genes in different tissues, developmental stage seedlings, stress-treated seedlings and all experimental groups combined. The average pairwise variations Vn/Vn+1 were analysed between the normalization factors NFn and NFn+1 to indicate the optimal number of reference genes in different samples.

NormFinder with lower values indicating higher stability, is another Excel application, which ranks the candidate genes from their minimal combined inter- and intra-group variation of expression.

Table 3 shows the expression stability values of the four data sets calculated by NormFinder. *ACT2, TUB-β* and *EF1-α* were the top three suitable genes for normalization, while ACT3 was at the end of the rankings in all groups. *ACT7* ranks at the top of expression stability in the ST group, but it is very unstable in other groups. The stability order of the candidate genes calculated by NormFinder is nearly the same as that calculated by GeNorm, but this conclusion did not exclude a small difference. For example, *ACT2* was identified as the most stable reference gene in the TS group with the GeNorm analysis, while its stability rankings placed it third within the NormFinder analysis.

**Table 3.** Ranking and expression stability values of the candidate reference genes calculated by NormFinder.

| Rank | Total |  | Tissue |  | Stage |  | Stress |  |
| --- | --- | --- | --- | --- | --- | --- | --- | --- |
|  | Gene | Stability Value | Gene | Stability Value | Gene | Stability Value | Gene | Stability Value |
| 1 | *TUB-β* | 0.246 | *EF1-α* | 0.090 | *ACT2* | 0.048 | *ACT2* | 0.141 |
| 2 | *ACT2* | 0.332 | *TUB-β* | 0.248 | *TUB-β* | 0.048 | *ACT7* | 0.179 |
| 3 | *CYP1* | 0.346 | *ACT2* | 0.323 | *TUB-α* | 0.190 | *EF1-α* | 0.194 |
| 4 | *EF1-α* | 0.398 | *CYP1* | 0.371 | *EF1-α* | 0.205 | *TUB* | 0.195 |
| 5 | *TUB-α* | 0.445 | *ACT7* | 0.462 | *CYP1* | 0.251 | *CYP1* | 0.205 |
| 6 | *SAND* | 0.453 | *UBC2* | 0.521 | *GSA* | 0.304 | *TUA* | 0.247 |
| 7 | *ACT7* | 0.511 | *GSA* | 0.620 | *GAPDH* | 0.308 | *GSA* | 0.262 |
| 8 | *UBC2* | 0.532 | *PP2A* | 0.644 | *PP2A* | 0.325 | *PKSI* | 0.311 |
| 9 | *PKSI* | 0.562 | *GAPDH* | 0.653 | *SAND* | 0.337 | *SAND* | 0.335 |
| 10 | *PP2A* | 0.586 | *PKSI* | 0.663 | *UBC2* | 0.487 | *UBC2* | 0.341 |
| 11 | *RPL13* | 0.594 | *SAND* | 0.687 | *RPL13* | 0.491 | *RPL13* | 0.353 |
| 12 | *GSA* | 0.602 | *TUB-α* | 0.710 | *ACT7* | 0.540 | *GAPDH* | 0.375 |
| 13 | *GAPDH* | 0.668 | *ROL13* | 0.789 | *PKSI* | 0.771 | *PP2A* | 0.401 |
| 14 | *ACT3* | 0.688 | *ACT3* | 0.934 | *ACT3* | 1.297 | *ACT3* | 0.462 |

BestKeeper is also an Excel-based program that is able to compare the expression levels of up to ten target genes, each in up to a hundred biological samples [33]. The CV and SD of the fourteen candidate genes computed by BestKeeper are represented in Table 4. The smaller the SD and the CV, the better the stability of the internal reference genes. When SD >1, the expression of the reference gene was unstable [33]. The results revealed that *ACT2* (SD = 0.303, CV = 1.121) and *TUB-β* (SD = 0.551, CV = 2.145) were the top two most stable genes for the TT samples. *TUB-β* was ranked first in the TS samples and PKS1 showed the lowest stability. *ACT2* was still the most stable gene in the SG and ST samples. Unlike the GeNorm and NormFinder rankings, the stability of *RPL13* was ahead in BestKeeper analysis and was relatively stable under all tested conditions.

**Table 4.** Ranking of the candidate reference genes and their expression stability calculated by BestKeeper.

| Rank | Total |  |  | Tissue |  |  | Stage |  |  | Stress |  |  |
| --- | --- | --- | --- | --- | --- | --- | --- | --- | --- | --- | --- | --- |
|  | Gene | SD[±CP] | CV | Gene | SD[±CP] | CV | Gene | SD[±CP] | CV | Gene | SD[±CP] | CV |
| 1 | *ACT2* | 0.303 | 1.121 | *TUB-β* | 0.184 | 0.677 | *TUB-β* | 0.294 | 0.996 | *ACT2* | 0.181 | 0.692 |
| 2 | *TUB-β* | 0.551 | 2.145 | *RPL13* | 0.520 | 2.106 | *ACT2* | 0.329 | 1.514 | *EF1-α* | 0.231 | 0.857 |
| 3 | *CYP1* | 0.597 | 2.308 | *CYP-1* | 0.766 | 2.963 | *GSA* | 0.454 | 2.020 | *CYP-1* | 0.274 | 1.070 |
| 4 | *TUB-α* | 0.606 | 2.650 | *TUB-α* | 0.798 | 3.291 | *RPL13* | 0.585 | 3.098 | *TUB-β* | 0.281 | 1.103 |
| 5 | *RPL13* | 0.610 | 2.423 | *SAND* | 0.870 | 3.819 | *GAPDH* | 0.610 | 2.258 | *RPL13* | 0.285 | 1.167 |
| 6 | *EF1-α* | 0.628 | 2.099 | *ACT7* | 0.925 | 3.640 | *EF1-α* | 0.610 | 2.378 | *PKS-1* | 0.319 | 1.678 |
| 7 | *ACT7* | 0.633 | 2.452 | *ACT2* | 0.943 | 3.641 | *SAND* | 0.695 | 2.771 | *TUB-α* | 0.356 | 1.499 |
| 8 | *PP2A* | 0.644 | 2.655 | *EF1-α* | 0.958 | 3.787 | *CYP1* | 0.793 | 3.036 | *SAND* | 0.357 | 1.550 |
| 9 | *SAND* | 0.763 | 3.248 | *GAPDH* | 0.976 | 4.580 | *TUB-α* | 0.803 | 3.540 | *ACT3* | 0.362 | 1.440 |
| 10 | *UBC2* | 0.770 | 4.005 | *ACT3* | 0.985 | 3.656 | *UBC2* | 0.833 | 3.211 | *PP2A* | 0.378 | 1.471 |
| 11 | *PKSI* | 0.831 | 3.498 | *PP2A* | 0.995 | 4.151 | *PKS-I* | 0.855 | 3.732 | *ACT7* | 0.399 | 1.705 |
| 12 | *GSA* | 0.893 | 3.928 | *GSA* | 1.013 | 5.071 | *PP2A* | 0.859 | 3.539 | *UBC2* | 0.436 | 1.457 |
| 13 | *GAPDH* | 0.947 | 4.803 | *UBC2* | 1.165 | 3.841 | *ACT7* | 0.873 | 4.605 | *GSA* | 0.471 | 2.090 |
| 14 | *ACT3* | 1.039 | 3.989 | *PKS-I* | 1.393 | 5.655 | *ACT3* | 1.888 | 7.201 | *GAPDH* | 0.562 | 2.909 |

### Reference gene validation

In order to test the reliability of the results, the relative expression patterns for the target gene *HpHYP1*, belonging to the PR-10 family [35,36] and associated with stress control, were evaluated by different internal control genes in the roots, stems, leaves and flowers. Through the present research results, the top two stable genes (*ACT2* and *TUB-β*) and the most unstable gene (*ACT3*) were used as internal controls. As shown in Fig 5, when normalized using *ACT2* as the reference genes, transcript abundance of *HpHYP1* was upregulated compared with the result in the root samples. When ACT2 and *TUB-β* (as identified by geNorm) both served as the internal references, the expression patterns are similiar with *ACT2*. When normalization was based on *TUB-β alone*, the expression level of *HpHYP1* was still up-regulation. The only difference was that the expression level in stems was lower than that in the leaves, but a significant difference was not apparent. In general, when stable reference genes including *ACT2, TUB-β* and their combination were used as internal parameters, *HpHYP1* had the highest expression level in flowers. When normalized by the less stable gene *ACT3*, the target genes in all tissues were expressed at lower levels but showed significantly upregulation (*P* < 0.001). Thus, selection of inappropriate reference genes can lead to over- or under-estimation of the relative transcript level, which might lead to a biased result.

**Figure 5.**
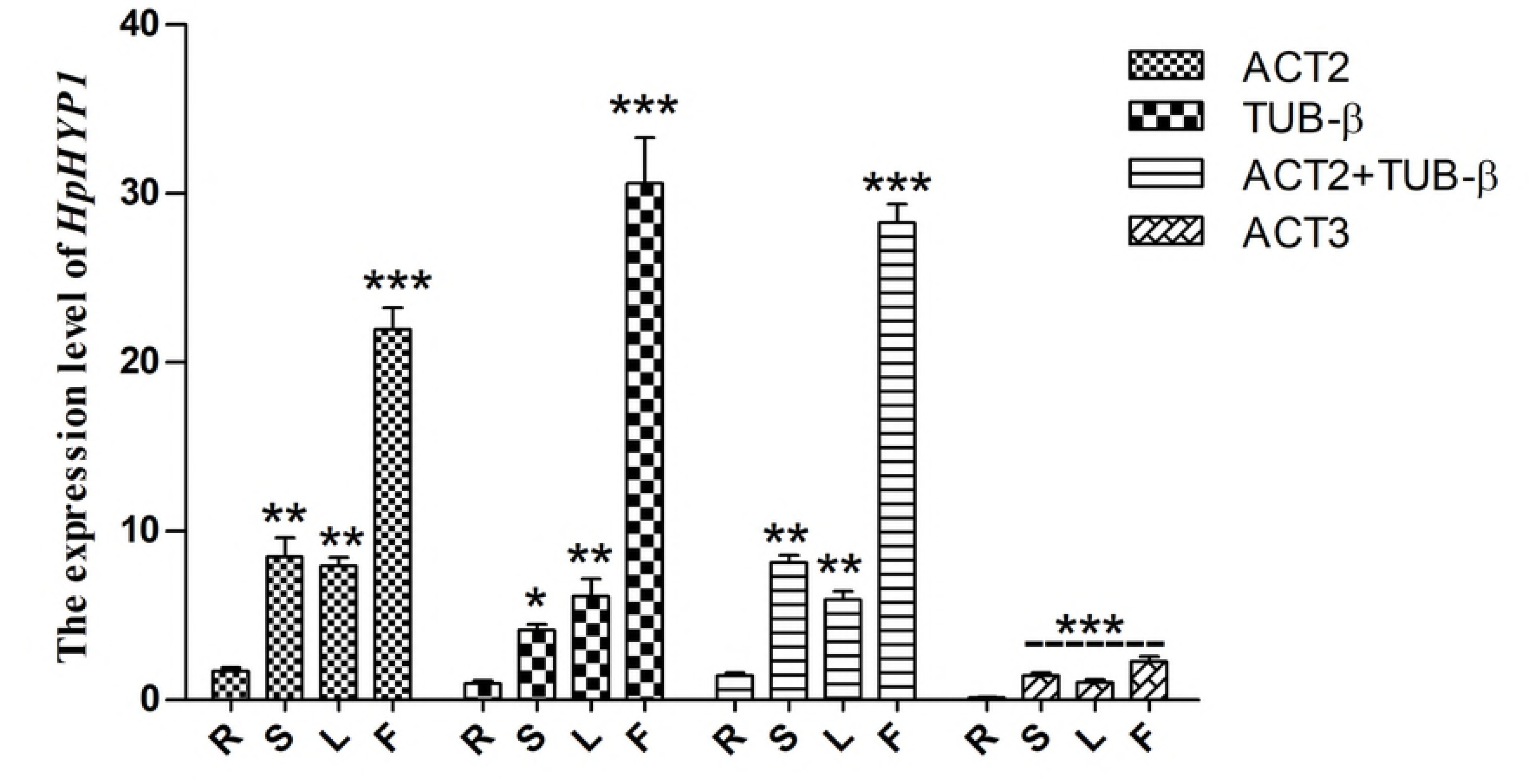
Relative expression levels of *HpHYP1* in different tissues (root, stem, leaf, and flower) used for normalization using the most stable reference genes or a combination and the least stable genes. Error bars show mean standard error calculated from three biological replicates. The statistical level was assessed with * P < 0.05, ** P < 0.01, and *** P < 0.001.

## Discussion

qRT-PCR is a very convenient and efficient method and widely used for gene expression analysis [8,15]. The reliability of qRT-PCR data will be greatly improved by the inclusion of a reference gene with a transcription level that is invariable across different experimental conditions [8]. In many published studies, some common reference genes especially the housekeeping genes have been verified in many species, such as in *Arabidopsis thaliana* [46,54], *Poa pratensis* [54], *Salvia miltiorrhiza* [55], *Cucumis sativus* [56], *Gentiana macrophylla* [37], and *Isatis indigotica* [57]. But selecting appropriate reference genes for *H. perforatums* in different experimental conditions have not yet been conducted, which might be due to the fact that genomic information for *H. perforatums* is very limited in NCBI. With the rapid development of whole-genome sequencing technology, deciphering the genomes of medicinal herbs is a vital step in understanding and improving their medicinal value. For that reason, we completed the assembly of the first high-quality sequence of the *H. perforatum* genome. Therefore, the reported literature on *H. perforatum* [58,59] combined withthe obtained genomic data laid the foundation for selecting reference genes.

Housekeeping genes like those participated in cell structure maintenance (*ACT* and *TUB*) or basic cellular processes (*UBC* and *CYP*) remain widely applied, but their expression can vary between different types of tissue (normal and pathological samples) and under different treatment conditions (drugs and chemicals) [60,61]. Thus, normalization with multiple reference genes is becoming popular and standard in plant research [33,53]. In this work, fourteen potential reference genes were chosen, including nine traditional housekeeping genes and five potential reference genes. The study represents an effort to identify and systematically compare fourteen potential reference genes. We assessed the expression patterns of these genes in different tissues, different developmental stages and under various abiotic stress treatments to screen for the most stable housekeeping genes for qRT-PCR analyses. As shown in Fig. 2, No one has a constant CT value, which shows how important it is to evaluate the most suitable reference gene for normalizing expression under all detection conditions in *H. perforatum*.

The analytical procedures applied in our research based on statistical algorithm to identify the stability of reference genes are commonly used by researchers to select the best reference genes [53,62–65]. Furthermore, a comparison of the different algorithms of reference genes contributes a better evaluation and reduces the risk of artificial selection of co-regulated transcripts [66]. The arithmetic of NormFinder and geNorm is almost the same, but GeNorm analysis is used to determine not only the most stable reference genes but also the optimal number of the gene combinations. In the present study, when the data were taken together to figure out the optimal number of reference genes, the pairwise variation of V2/3 values for all of the experimental sets was lower than the cut off threshold of 0.15 (Fig. 4). Thus, these results show that the best combination (*ACT2* and *TUB-β*) should be both involved to improve the accuracy of the quantitative expression analysis in *H. perforatum*. For the BestKeeper program, it can worked out the best suited standards in accordance with SD and CV values [33], out of ten candidates including reference and target genes.

In our research, GeNorm and NormFinder create similar rankings for stability values, while BestKeeper programs always get different rankings. For example, BestKeeper ranked RPL13 as the most stable, as it was relatively stable under all tested conditions. In contrast, the expression stability value of the RPL13 gene in GeNorm and Normfinder was very low. There were also some previous papers revealing similar difference between BestKeeper and other methods [54,67]. Homologous genes are widely used as reference genes for gene expression analysis. For example, *ACT6, ACT8,* and *ACT7* were selected as internal controls for stress treatments in *Fortunella crassifolia* [33]; *UBC19, UBC22*, and *UBC29* were selected as reference genes in the context of the relevant experimental conditions in *Isatis indigotica* [57]; and *EF1A2a, EF1A2b*, and *EF1A1a1* were the best reference genes under all tested conditions in *Glycine max* [68]. Nevertheless, as can be seen in our results, the three homologous genes (*ACT2, ACT3* and *ACT7*) exhibited totally different expression levels when used for the normalization of qRT-PCR, especially *ACT2* and *ACT3* (Table 2, Table 3, Table 4 and Fig. 3). Therefore the paralogous genes have similar stuctures, but their expression levels are entirely differnet in gene expression quantification [57].

In summary, our results support that the selection of reference genes has a significant impact on the normalized gene expression data in qRT-PCR experiments. We have investigated the expression of fourteen candidate reference genes across a large number of *H. perforatum* samples in an effort to identify those most stable genes for normalizing gene expression. Finally, a combination of the two genes *ACT2* and *TUB-β* provided the best internal controls for transcript normalization across experimental conditions in this study.

## Acknowledgements

This work was supported by the National Natural Science Foundation of China (31670299) and the Fundamental Research Funds for the Central Universities (1301031470, GK201806006).

## Author contributions

WZ managed and carried out the experimental design. LY and YS sowed the seeds, directed the abiotic stress treatments and collected samples. WZ and LY screened the candidate reference genes and designed the primers. QZ, BW and LL performed the qRT-PCR. WZ drafted the manuscript, and BL revised it. All authors discussed and commented on the manuscript.

## Reference

1. Morenorisueno MA, Norman JMV, Moreno A, Zhang J, Ahnert SE, et al. (2010) Oscillating Gene Expression Determines Competence for Periodic Arabidopsis Root Branching. Science 329: 1306–1311.

2. Hirotaka Y, Hiroyuki F, Tomohito A, Akio O, Tsukasa N, et al. (2010) Gene expression analysis in cadmium-stressed roots of a low cadmium-accumulating solanaceous plant, Solanum torvum. Journal of Experimental Botany 61: 423–437.

3. Di W, Pan Y, Zhao X, Zhu L, Fu B, et al. (2011) Genome-wide temporal-spatial gene expression profiling of drought responsiveness in rice. Bmc Genomics 12: 149.

4. Dechorgnat J, Patrit O, Krapp A, Fagard M, Danielvedele F (2012) Characterization of the Nrt2.6 Gene in Arabidopsis thaliana: A Link with Plant Response to Biotic and Abiotic Stress. Plos One 7: e42491.

5. Bustin SA (2002) Bustin, S. A. Quantification of mRNA using real-time reverse transcription PCR (RT-PCR): trends and problems. J Mol Endocrinol. Journal of Molecular Endocrinology 29: 23–39.

6. Bustin SA, Benes V, Nolan T, Pfaffl MW (2005) Quantitative real-time RT-PCR – a perspective. Journal of Molecular Endocrinology 34: 597–601.

7. Nolan T, Hands RE, Bustin SA (2006) Quantification of mRNA using real-time RT-PCR. Nature Protocols 1: 1559.

8. Huggett J, Dheda K, Bustin S, Zumla A (2005) Real-time RT-PCR normalisation; strategies and considerations. Genes & Immunity 6: 279–284.

9. Udvardi MK, Czechowski T, Scheible WR (2008) Eleven golden rules of quantitative RT-PCR. Plant Cell 20: 1736–1737.

10. Bustin SA, Mueller R (2005) Real-time reverse transcription PCR (qRT-PCR) and its potential use in clinical diagnosis. Clinical Science 109: 365–379.

11. Leal MF, Astur DC, Debieux P, Arliani GG, Silveira Franciozi CE, et al. (2015) Identification of Suitable Reference Genes for Investigating Gene Expression in Anterior Cruciate Ligament Injury by Using Reverse Transcription-Quantitative PCR. Plos One 10: e0133323.

12. He Y, Yan H, Hua W, Huang Y, Wang Z (2016) Selection and Validation of Reference Genes for Quantitative Real-time PCR in Gentiana macrophylla. Frontiers in Plant Science 7: 945.

13. Tong ZG, Gao ZH, Wang F, Zhou J, Zhang Z (2009) Selection of reliable reference genes for gene expression studies in peach using real-time PCR. BMC Molecular Biology,10,1(2009-07- 20) 10: 1–13.

14. Gutierrez L, Mauriat M, Guénin S, Pelloux J, Lefebvre JF, et al. (2010) The lack of a systematic validation of reference genes: a serious pitfall undervalued in reverse transcription-polymerase chain reaction (RT-PCR) analysis in plants. Plant Biotechnology Journal 6: 609–618.

15. Brattelid T, Levy FO (2011) Quantification of GPCR mRNA Using Real-Time RT-PCR: Humana Press. 165–193 p.

16. Bustin SA, Nolan T (2004) Pitfalls of quantitative real-time reverse-transcription polymerase chain reaction. J Biomol Tech 15: 155–166.

17. Butterweck V (2003) Mechanism of action of St John’s wort in depression : what is known? Cns Drugs 17: 539.

18. Birt DF, Widrlechner MP, Hammer KDP, Hillwig ML, Wei JQ, et al. (2009) Hypericum in infection: identification of anti-viral and anti-inflammatory constituents. Pharmaceutical Biology 47: 774–782.

19. Caraci F, Crupi R, Drago F, Spina E (2011) Metabolic drug interactions between antidepressants and anticancer drugs: focus on selective serotonin reuptake inhibitors and hypericum extract. Current Drug Metabolism 12: –.

20. Schröder J (1997) A family of plant-specific polyketide synthases: facts and predictions. Trends in Plant Science 2: 373–378.

21. Velada I, Ragonezi C, Arnholdt-Schmitt B, Cardoso H (1932) Reference Genes Selection and Normalization of Oxidative Stress Responsive Genes upon Different Temperature Stress Conditions in Hypericum perforatum L. Plos One 9: e115206.

22. Willems E, Leyns L, Vandesompele J (2008) Standardization of real-time PCR gene expression data from independent biological replicates. Analytical Biochemistry 379: 127–129.

23. Imai T, Ubi BE, Saito T, Moriguchi T (2014) Evaluation of reference genes for accurate normalization of gene expression for real time-quantitative PCR in Pyrus pyrifolia using different tissue samples and seasonal conditions. Plos One 9: e86492.

24. Llanos A, François JM, Parrou JL (2015) Tracking the best reference genes for RT-qPCR data normalization in filamentous fungi. BMC Genomics,16,1(2015-02-14) 16: 71.

25. Goulao LF, Fortunato AS, Ramalho JC (2012) Selection of Reference Genes for Normalizing Quantitative Real-Time PCR Gene Expression Data with Multiple Variables in Coffea spp. Plant Molecular Biology Reporter 30: 741–759.

26. Costa MD, Duro N, Batista-Santos P, Ramalho JC, Ribeiro-Barros AI (2015) Validation of candidate reference genes for qRT-PCR studies in symbiotic and non-symbiotic Casuarina glauca Sieb. ex Spreng. under salinity conditions. Symbiosis 66: 21–35.

27. Thellin O, Zorzi W, Lakaye B, Borman BD, Coumans B, et al. (1999) Housekeeping genes as internal standards: use and limits. Journal of Biotechnology 75: 291–295.

28. Selvey S, Thompson EW, Matthaei K, Lea RA, Irving MG, et al. (2001) Beta-actin–an unsuitable internal control for RT-PCR. Molecular & Cellular Probes 15: 307–311.

29. Ohl F, Jung M, Xu C, Stephan C, Rabien A, et al. (2005) Gene expression studies in prostate cancer tissue: which reference gene should be selected for normalization? Journal of Molecular Medicine 83: 1014–1024.

30. Vandesompele J, De KP, Pattyn F, Poppe B, Van NR, et al. (2002) Accurate normalization of real-time quantitative RT-PCR data by geometric averaging of multiple internal control genes. Genome Biology 3: research0034.0031.

31. Kidd M, Nadler B, Mane S, Eick G, Malfertheiner M, et al. (2007) GeneChip, geNorm, and gastrointestinal tumors: novel reference genes for real-time PCR. Physiological Genomics 30: 363–370.

32. Wang Q, Ishikawa T, Michiue T, Zhu BL, Guan DW, et al. (2012) Stability of endogenous reference genes in postmortem human brains for normalization of quantitative real-time PCR data: comprehensive evaluation using geNorm, NormFinder, and BestKeeper. International Journal of Legal Medicine 126: 943–952.

33. Pfaffl MW, Tichopad A, Prgomet C, Neuvians TP (2004) Determination of stable housekeeping genes, differentially regulated target genes and sample integrity: BestKeeper–Excel-based tool using pair-wise correlations. Biotechnology Letters 26: 509–515.

34. Li T, Wang J, Lu M, Zhang T, Qu X, et al. (2017) Selection and Validation of Appropriate Reference Genes for qRT-PCR Analysis inIsatis indigoticaFort. Frontiers in Plant Science 8.

35. Košuth J, Katkovčinová Z, Olexová P, Čellárová E (2007) Expression of the hyp-1 gene in early stages of development of Hypericum perforatum L. Plant Cell Reports 26: 211.

36. Assavalapsakul W, Panyim S (2012) Molecular cloning and tissue distribution of the Toll receptor in the black tiger shrimp, Penaeus monodon. Genetics & Molecular Research Gmr 11: 484.

37. Radonić A, Thulke S, Mackay IM, Landt O, Siegert W, et al. (2004) Guideline to reference gene selection for quantitative real-time PCR. Biochemical & Biophysical Research Communications 313: 856–862.

38. Schlotter YM, Veenhof EZ, Brinkhof B, Rutten VP, Spee B, et al. (2009) A GeNorm algorithm-based selection of reference genes for quantitative real-time PCR in skin biopsies of healthy dogs and dogs with atopic dermatitis. Veterinary Immunology & Immunopathology 129: 115–118.

39. Zhang X, Xu ZC, Xu J, Ji AJ, Luo HM, et al. (2016) Selection and validation of reference genes for normalization of quantitative real-time reverse transcription PCR analysis in Poria cocos (Schw.) Wolf (Fuling). Chinese Medicine 11: 1–17.

40. Die JV, Roman B, Obrero A (2011) Selection of Reference Genes for Gene Expression Studies in Zucchini (Cucurbita pepo) Using qPCR. J Agric Food Chem 59: 5402–5411.

41. Rhinn H, Marchand-Leroux C, Croci N, Plotkine M, Scherman D, et al. (2008) Housekeeping while brain’s storming Validation of normalizing factors for gene expression studies in a murine model of traumatic brain injury. Bmc Molecular Biology 9: 62–62.

42. Rotenberg D, Thompson TS, German TL, Willis DK (2006) Methods for effective real-time RT-PCR analysis of virus-induced gene silencing. Journal of Virological Methods 138: 49–59.

43. Zhang Q, Li H, Fan C, Hu R, Fu YF (2009) Evaluation of putative reference genes for gene expression normalization in soybean by quantitative real-time RT-PCR. BMC Molecular Biology,10,1(2009-09-28) 10: 93–93.

44. Tong Z, Gao Z, Fei W, Zhou J, Zhen Z (2009) Selection of reliable reference genes for gene expression studies in peach using real-time PCR. Bmc Molecular Biology 10: 1–13.

45. Guo Y, Chen JX, Yang S, Fu XP, Zhang Z, et al. (2010) Selection of reliable reference genes for gene expression study in nasopharyngeal carcinoma. 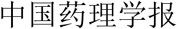 31: 1487–1494.

46. Dekkers BJ, Willems L, Bassel GW, van BolderenVeldkamp RP, Ligterink W, et al. (2012) Identification of reference genes for RT-qPCR expression analysis in Arabidopsis and tomato seeds. Plant & Cell Physiology 53: 28–37.

47. Marchal E, Hult EF, Huang J, Tobe SS (2013) Sequencing and validation of housekeeping genes for quantitative real-time PCR during the gonadotrophic cycle of Diploptera punctata. BMC Research Notes,6,1(2013-06-19) 6: 1–10.

48. Yang Z, Chen Y, Hu B, Tan Z, Huang B (2015) Identification and Validation of Reference Genes for Quantification of Target Gene Expression with Quantitative Real-time PCR for Tall Fescue under Four Abiotic Stresses. Plos One 10: e0119569.

49. Migocka M, Papierniak A (2011) Identification of suitable reference genes for studying gene expression in cucumber plants subjected to abiotic stress and growth regulators. Molecular Breeding 28: 343–357.

50. Swift ML (1997) GraphPad Prism, Data Analysis, and Scientific Graphing. Journal of Chemical Information & Modeling 37: 411–412.

51. Michalska K, Fernandes H, Sikorski M, Jaskolski M (2010) Crystal structure of Hyp-1, a St. John’s wort protein implicated in the biosynthesis of hypericin. Journal of Structural Biology 169: 161–171.

52. Kumar V, Sharma R, Trivedi PC, Vyas GK, Khandelwal V (2011) Traditional and novel references towards systematic normalization of qRT-PCR data in plants. Australian Journal of Crop Science 5: 1455–1468.

53. Vandesompele J, Preter KD, Pattyn F, Poppe B, Roy NV, et al. (2002) Accurate normalization of real-time quantitative RT-PCR data by geometric averaging of multiple internal control genes. Genome Biology 3: research0034.0031.

54. Czechowski T, Stitt M, Altmann T, Udvardi MK, Scheible WR (2005) Genome-wide identification and testing of superior reference genes for transcript normalization in Arabidopsis. Plant Physiology 139: 5–17.

55. Espinosa JR, Ayub Q, Chen Y, Xue Y, Tylersmith C (2015) Structural variation on the human Y chromosome from population-scale resequencing. Croatian Medical Journal 56: 194–207.

56. Yi S, Qian Y, Han L, Sun Z, Fan C, et al. (2012) Selection of reliable reference genes for gene expression studies in Rhododendron micranthum Turcz. Scientia Horticulturae 138: 128–133.

57. Li T, Wang J, Lu M, Zhang T, Qu X, et al. (2017) Selection and Validation of Appropriate Reference Genes for qRT-PCR Analysis in Isatis indigotica Fort. Frontiers in Plant Science 8.

58. Yao L (2011) Molecular cloning and tissue-specific expression of two different chitin deacetylase cDNA sequences from Mamestra brassicae. Chinese Journal of Applied Entomology 48: 1417–1424.

59. Velada I, Ragonezi C, Arnholdtschmitt B, Cardoso H (1932) Reference Genes Selection and Normalization of Oxidative Stress Responsive Genes upon Different Temperature Stress Conditions in Hypericum perforatum L. Plos One 9: e115206.

60. Thorrez L, Deun KV, Tranchevent L, Lommel LV, Engelen K, et al. (2008) Using Ribosomal Protein Genes as Reference: A Tale of Caution. Plos One 3: e1854.

61. Robinson TL, Sutherland IA, Sutherland J (2007) Validation of candidate bovine reference genes for use with real-time PCR. Veterinary Immunology & Immunopathology 115: 160–165.

62. Haller F, Kulle B, Schwager S, Gunawan B, Von HA, et al. (2004) Equivalence test in quantitative reverse transcription polymerase chain reaction: confirmation of reference genes suitable for normalization. Analytical Biochemistry 335: 1–9.

63. Jarošová J, Kundu JK (2010) Validation of reference genes as internal control for studying viral infections in cereals by quantitative real-time RT-PCR. Bmc Plant Biology 10: 146–146.

64. Szabo A, Perou CM, Karaca M, Perreard L, Quackenbush JF, et al. (2008) Statistical modeling for selecting housekeeper genes. Genome Biology 9: 405–405.

65. Kok JBD, Roelofs RW, Giesendorf BA, Pennings JL, Waas ET, et al. (2005) Normalization of gene expression measurements in tumor tissues: comparison of 13 endogenous control genes. Laboratory investigation; a journal of technical methods and pathology 85: 154.

66. Ayers D, Clements DN, Salway F, Day PJ (2007) Expression stability of commonly used reference genes in canine articular connective tissues. Bmc Veterinary Research 3: 1–10.

67. Rapacz M, Stępień A, Skorupa K (2012) Internal standards for quantitative RT-PCR studies of gene expression under drought treatment in barley (Hordeum vulgare L.): the effects of developmental stage and leaf age. Acta Physiologiae Plantarum 34: 1723–1733.

68. Bansal R, Mittapelly P, Cassone BJ, Mamidala P, Redinbaugh MG, et al. (2015) Recommended Reference Genes for Quantitative PCR Analysis in Soybean Have Variable Stabilities during Diverse Biotic Stresses. Plos One 10: e0134890.

